## Supplementary figures and images for "CRB3 navigates Rab11 trafficking vesicles to promote γTuRC assembly during ciliogenesis"

### Figure 1-figure supplementary 1

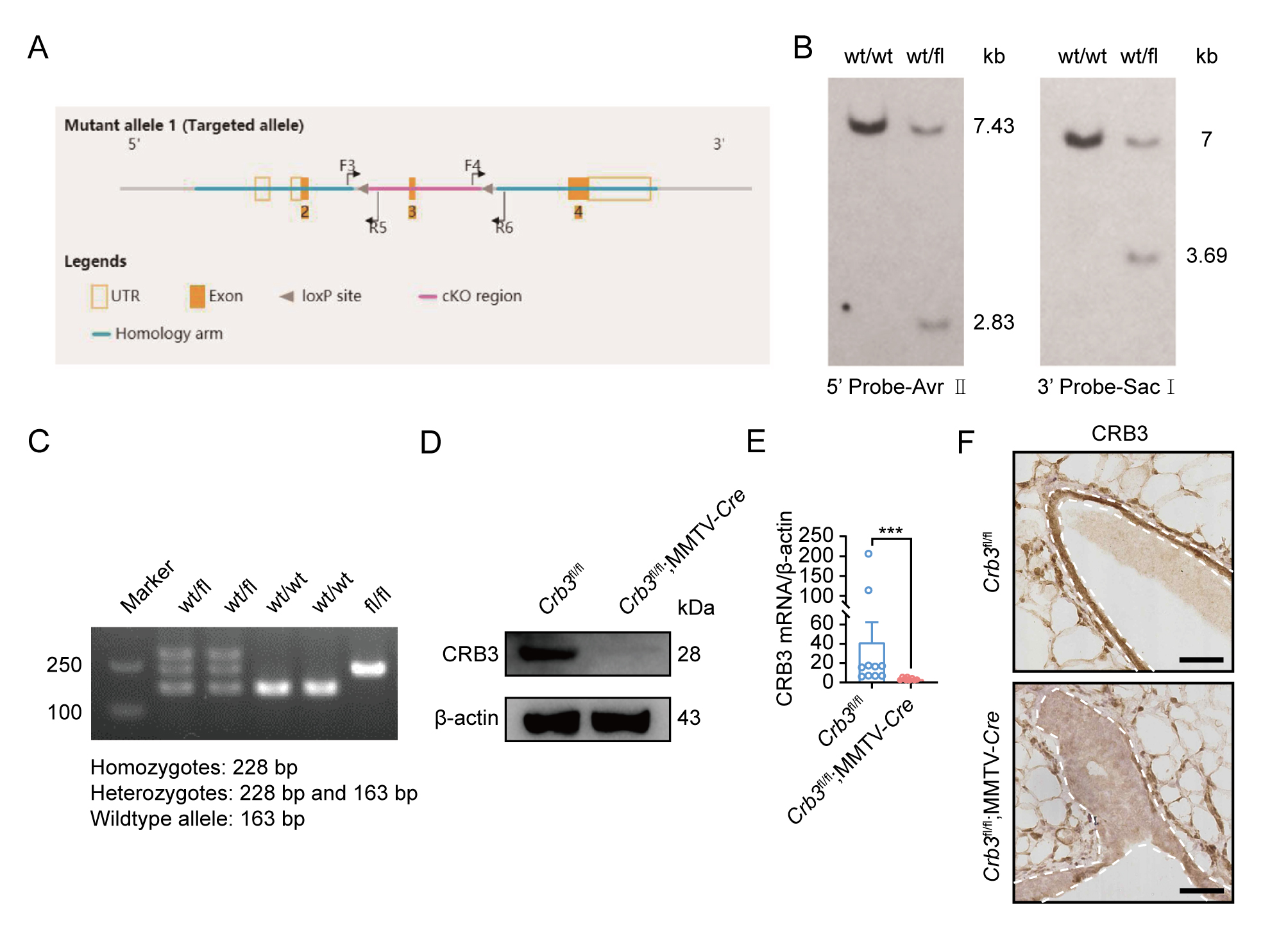

### Figure 2-figure supplementary 1

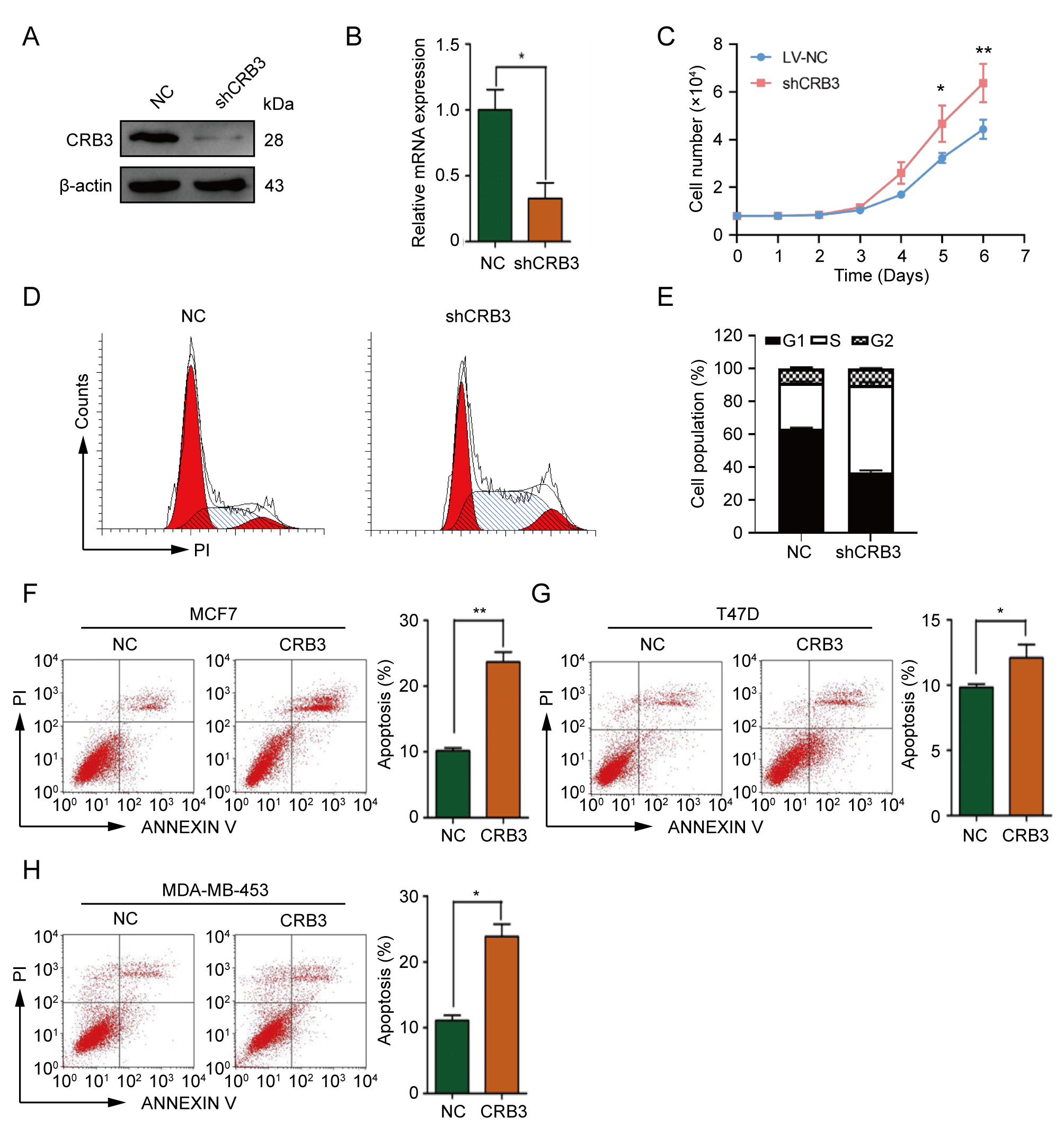

### Figure 3-figure supplementary 1

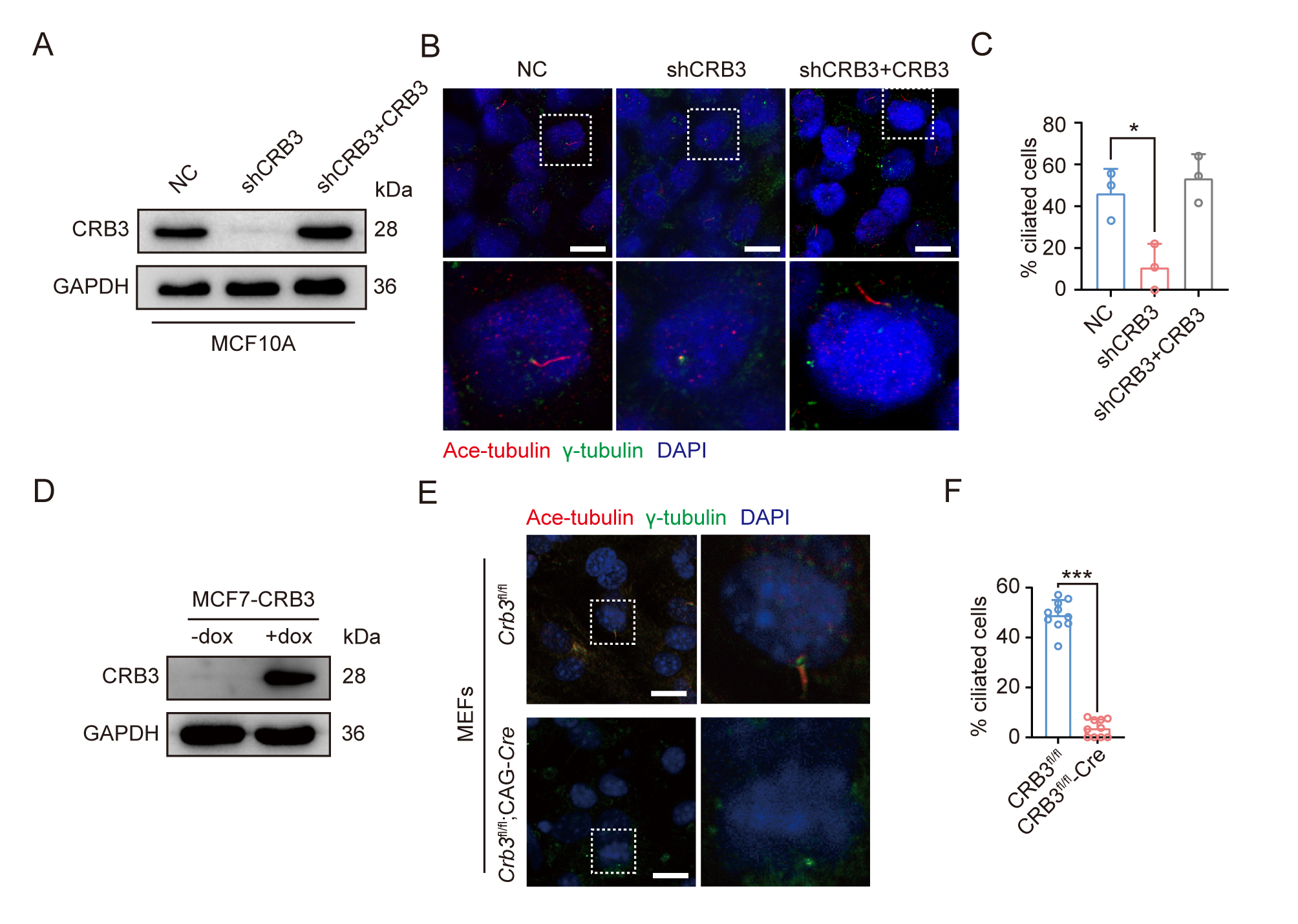

### Figure 5-figure supplementary 1

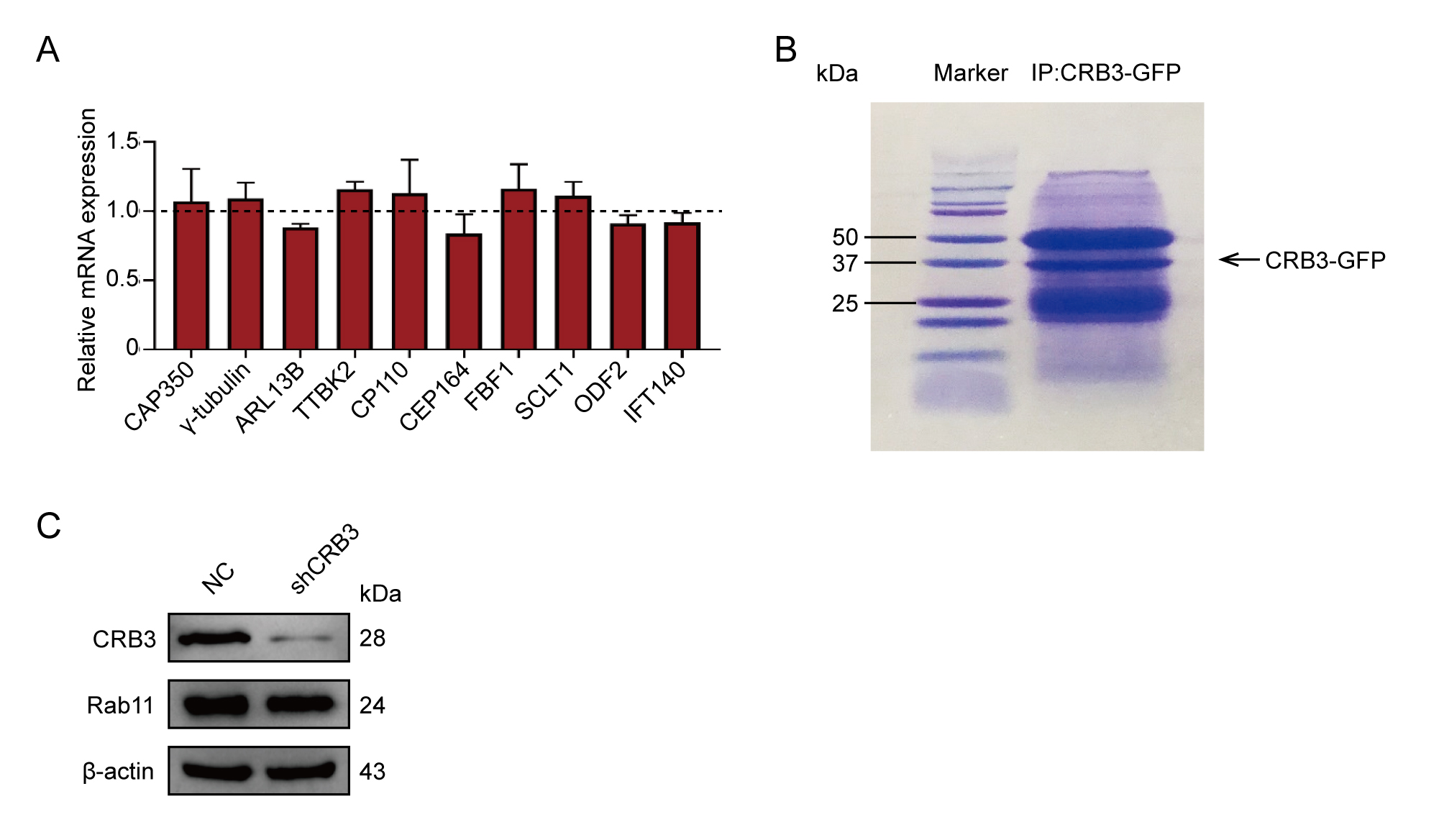

### Figure 5-figure supplementary 2

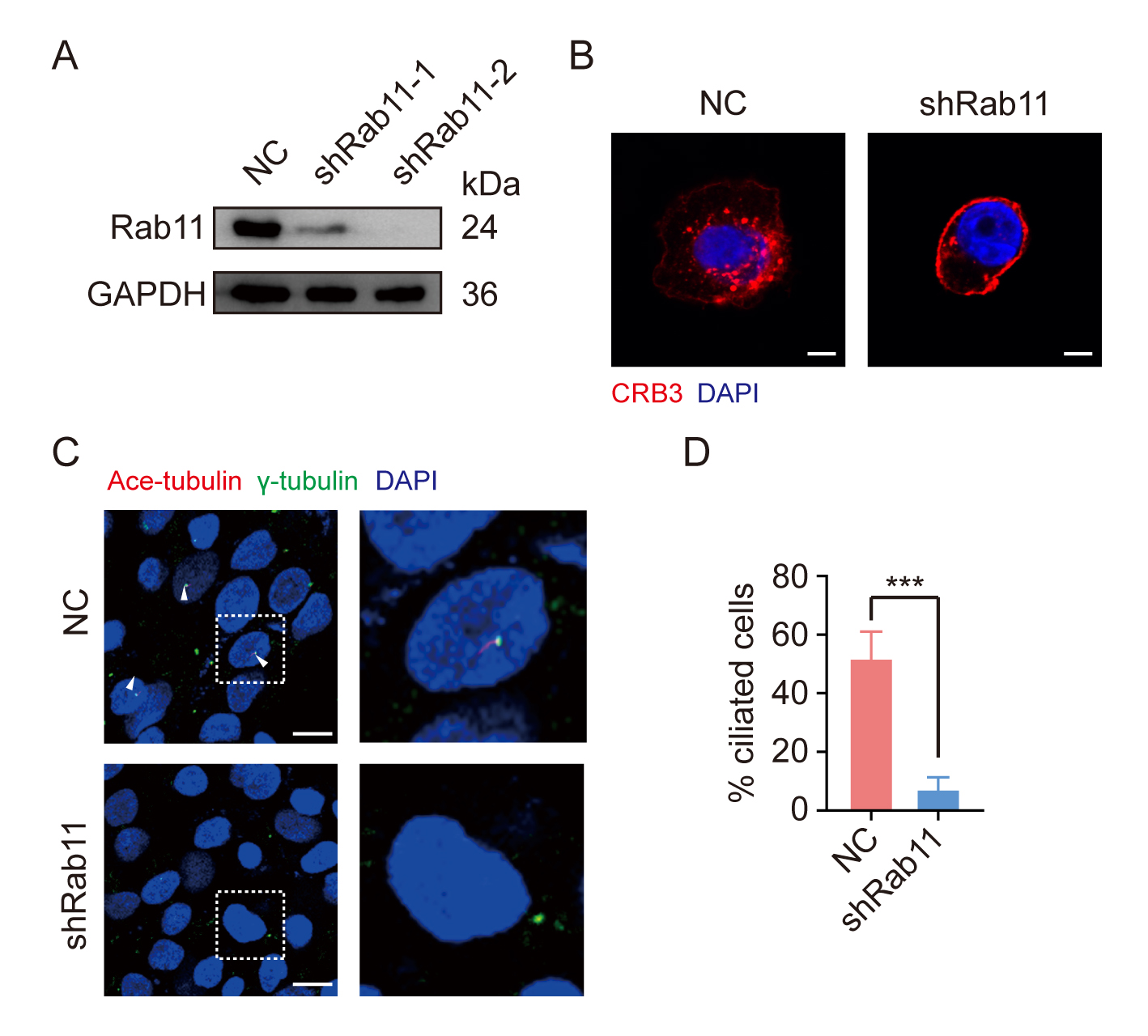

### Figure 8-figure supplementary 1

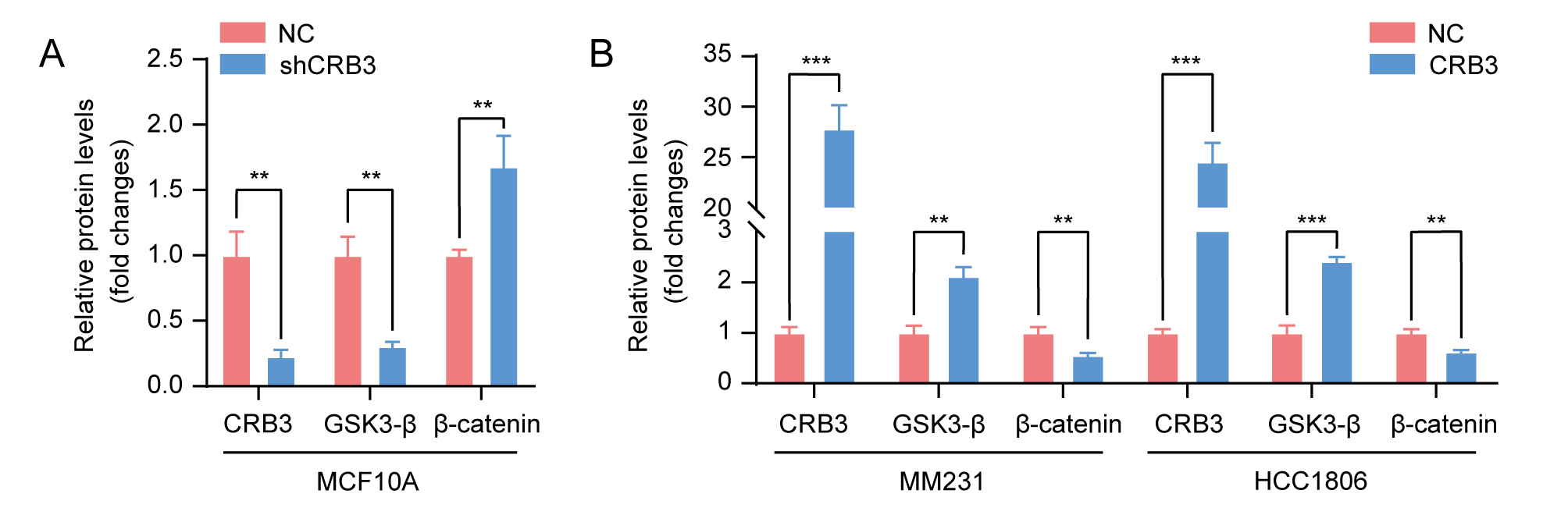
