## Supplementary File 1 for "CRB3 navigates Rab11 trafficking vesicles to promote γTuRC assembly during ciliogenesis"

**Supplementary File 1.** The sequences of primer pairs used in real-time PCR

| **Gene** |  | **Sequence (5’-3-)** |
| --- | --- | --- |
| *CRB3* | F | CTTCTGCAAATGAGAATAGCACTG |
|  | R | GAAGACCACGATGATAGCAGTGA |
| *Crb3* | F | CACCGGACCCTTTCACAAATA |
|  | R | CCCACTGCTATAAGGAGGACT |
| *CEP350* (CAP350) | F | ATCGTGTGGAATTTCGTGAACC |
|  | R | TCCGTTCTTCTCGACTGCCTA |
| *TUBG1* (γ-tubulin) | F | AGCTGGTGTCTACCATCATGT |
|  | R | CGTAGTGAGAGGGGTGTAGC |
| *ARL13B* | F | ATGTTCAGCAATCTCGGGGTA |
|  | R | TGTCTCTTTTTGGATGCGTTCAT |
| *TTBK2* | F | CAATCAACGCACATCGGAACA |
|  | R | GAGCCTACTTGCTCCTTGTCC |
| *CCP110* (CP110) | F | AGACGCAGTCTGAGAGGTAGT |
|  | R | CAGTGTTTGCCTGTCAACTGG |
| *CEP164* | F | GTTCCTCATTAGCCCCAGTTC |
|  | R | AGGCTCACGCTTTGAGATCC |
| *FBF1* | F | TCAGAACAGGAGGTTTTCCTCT |
|  | R | GGGGCTCTTGTCAGAAGCTG |
| *SCLT1* | F | AGCCAAATTAGAGCTGAGAGTTG |
|  | R | CAGACACCACATCCTTCTCCT |
| *ODF2* | F | TGGAGGCGGAAATGGATGG |
|  | R | CCTTGTCAGGGTGTTGATGTC |
| *IFT140* | F | TCCATTCTTGGCAGTTGCTTAC |
|  | R | CTCTCGACGTGTGTATCTGGC |
| *RAB11A* (RAB11) | F | CAACAAGAAGCATCCAGGTTGA |
|  | R | GCACCTACAGCTCCACGATAAT |
| *GLI1* | F | AGCGTGAGCCTGAATCTGTG |
|  | R | CAGCATGTACTGGGCTTTGAA |
| *GAPDH* | F | CTGGGCTACACTGAGCACC |
|  | R | AAGTGGTCGTTGAGGGCAATG |
| *Gapdh* | F | AGGTCGGTGTGAACGGATTTG |
|  | R | TGTAGACCATGTAGTTGAGGTCA |
