## Supplementary File 2 for "CRB3 navigates Rab11 trafficking vesicles to promote γTuRC assembly during ciliogenesis"

**Supplementary File 2.** Characteristics of patients

| **No.** | **Age (Years)** | **Sex** | **Gender** | **Ethnicity** |
| --- | --- | --- | --- | --- |
| 1 | 71 | Female | Woman | Han/Chinese |
| 2 | 59 | Female | Woman | Han/Chinese |
| 3 | 46 | Female | Woman | Han/Chinese |
| 4 | 68 | Female | Woman | Han/Chinese |
| 5 | 55 | Female | Woman | Han/Chinese |
| 6 | 78 | Female | Woman | Han/Chinese |
| 7 | 51 | Female | Woman | Han/Chinese |
| 8 | 51 | Female | Woman | Han/Chinese |
| 9 | 52 | Female | Woman | Han/Chinese |
| 10 | 44 | Female | Woman | Han/Chinese |
| 11 | 58 | Female | Woman | Han/Chinese |
| 12 | 71 | Female | Woman | Han/Chinese |
| 13 | 47 | Female | Woman | Han/Chinese |
| 14 | 73 | Female | Woman | Han/Chinese |
| 15 | 54 | Female | Woman | Han/Chinese |
| 16 | 69 | Female | Woman | Han/Chinese |
| 17 | 78 | Female | Woman | Han/Chinese |
| 18 | 62 | Female | Woman | Han/Chinese |
| 19 | 78 | Female | Woman | Han/Chinese |
| 20 | 54 | Female | Woman | Han/Chinese |
| 21 | 42 | Female | Woman | Han/Chinese |
| 22 | 45 | Female | Woman | Han/Chinese |
| 23 | 45 | Female | Woman | Han/Chinese |
| 24 | 57 | Female | Woman | Han/Chinese |
| 25 | 53 | Female | Woman | Han/Chinese |
| 26 | 67 | Female | Woman | Han/Chinese |
| 27 | 51 | Female | Woman | Han/Chinese |
| 28 | 46 | Female | Woman | Han/Chinese |
| 29 | 49 | Female | Woman | Han/Chinese |
| 30 | 78 | Female | Woman | Han/Chinese |
| 31 | 81 | Female | Woman | Han/Chinese |
| 32 | 58 | Female | Woman | Han/Chinese |
| 33 | 52 | Female | Woman | Han/Chinese |
| 34 | 61 | Female | Woman | Han/Chinese |
| 35 | 47 | Female | Woman | Han/Chinese |
| 36 | 45 | Female | Woman | Han/Chinese |
| 37 | 48 | Female | Woman | Han/Chinese |
| 38 | 63 | Female | Woman | Han/Chinese |
| 39 | 39 | Female | Woman | Han/Chinese |
| 40 | 57 | Female | Woman | Han/Chinese |
| 41 | 47 | Female | Woman | Han/Chinese |
| 42 | 44 | Female | Woman | Han/Chinese |
| 43 | 44 | Female | Woman | Han/Chinese |
| 44 | 37 | Female | Woman | Han/Chinese |
| 45 | 62 | Female | Woman | Han/Chinese |
| 46 | 46 | Female | Woman | Han/Chinese |
| 47 | 53 | Female | Woman | Han/Chinese |
| 48 | 54 | Female | Woman | Han/Chinese |
| 49 | 46 | Female | Woman | Han/Chinese |
| 50 | 46 | Female | Woman | Han/Chinese |
